# Development of a DSM test battery to determine depression-like marmoset

**DOI:** 10.1101/2024.07.08.602574

**Authors:** Hajime Yamanaka, Hidetoshi Ishibashi, Katsuki Nakamura

## Abstract

Depression is a serious and high-incidence mental disorder that can lead to suicide. Despite much depression research using humans and rodents to date, it has been difficult to completely overcome depression and elucidate its etiology. Marmoset monkeys have been utilized to study neuropsychiatric disorders in recent years. Since the clinical diagnostic criteria for psychiatric disorders are mainly based on multiple behavioral changes, test battery systems, a multifaceted and comprehensive strategy, are applied to depression research. Here, we devised a test battery with six tests to investigate depression-like symptoms corresponding to clinical criteria and set criteria for a test indicator-based judgment. We did a usability confirmation experiment using a drug and judged three of nine marmosets as depression-like. This rate is comparable with the incidence rate of human patients; i.e., 10-30% of patients who received reserpine treatment suffer from depression. Thus, we propose the test battery we constructed in this study will contribute to the study of depression.

## Introduction

Major depressive disorder (depression) is a disease with numerous problems and in need of further research. Its lifetime prevalence is as high as 8-12%, and it affects about 280 million people worldwide^1–3^. The social and economic losses caused by depression are enormous, since depression can lead to suicide, social withdrawal, and absence from work. The World Health Organization has warned that depression will be one of the top risk diseases of global burden in the future, and the Lancet-World Psychiatric Association Committee has issued a declaration calling for efforts to reduce the global burden of depression^4,5^. The Diagnostic and Statistical Manual of Mental Disorders (DSM) diagnostic criteria for depression lists nine symptoms, implying that this disease exhibits a complex pathological condition in which multiple symptoms overlap at the same time^6^. Depression is a disease that is elusive and offers few clues, because not only has no clear etiology been identified, no clear abnormal findings in clinical tests, such as imaging or biomarkers, or postmortem brain examinations have been identified. Antidepressants, which are typical treatment, take two weeks to work, and one-third of patients are refractory to treatment^7,8^. Currently, patients continue to suffer from complex and long-term depressive symptoms, thereby prompting a vigorous pursuit of etiological elucidation within the scientific community. However, it is extremely difficult to find an etiology that goes beyond the realm of hypothesis. Given that depression has not been completely controlled or eradicated to date, expectations for future progress in depression research are high.

Depression studies typically use rodent models or clinical specimens; however, monkey models are beginning to be realized. Numerous rodent models based on drug and genetic manipulation techniques are reported^9,10^. Although the development of antidepressants using rodents has contributed significantly to depression research, no drug has been developed that can completely control depression. The genetic analysis of big data of human specimens has sought genetic predispositions and include large-scale genome-wide association studies (GWAS) that have reported associations with many genome sequences^11^. The reported etiological factors are being studied for treatment and diagnosis. On the other hand, monkey studies with depression are gradually beginning to progress, especially in model. Marmosets share many characteristics with humans and have been used extensively to study many human diseases but rarely depression^12–17^. One of the reasons is the lack of a well-developed research platform. Such a platform would provide its own unique insights while bridging rodent and human studies.

We devised the “DSM Test Battery for Depressed Monkeys” in this study in accordance with the policies and methods of the DSM, the international diagnostic standard for mental disorders. The DSM provides specific symptoms and diagnostic criteria and operationally makes a diagnosis according to them (symptom-based diagnosis). On the other hand, our test battery provides specific test indicators and criteria, and judgments of depression-like marmosets are made operationally according to them (test indicator-based judgment). Test indicators cover as many DSM depressive symptoms as possible, and model judgment is conducted on an individual basis. The Peephole Test (PhT) for visual exploratory behavior (VEB) and the Sucrose Preference Test (SPT) for anhedonic behavior were devised to examine “loss of interest and pleasure” in the DSM. The Eat Style (ES) test for food sorting behavior was devised to examine “fatigue or loss of energy” in the DSM. We added three tests corresponding to DSM depressive symptoms for a total of six tests developed. Some of these tests were developed based on the behavioral characteristics of monkeys and previous studies of depression. VEB dysfunction and anhedonic behavior closely mimic the loss of interest and pleasure, a core symptom of major depression episodes according to the DSM. VEB is spontaneous behavior exploring an external environment using vision and was first examined in rhesus monkey^18^. The same group later showed that the frequency of VEB increases when another monkey or a moving toy train is present and that the duration of VEB is prolonged when moving pictures are shown, indicating that exploratory behavior intensity depends on the type of stimulus^19,20^. Thus, it is argued that VEB is closely related to curiosity, interest, and motivation^21^. However, the behavioral characteristics of other species including marmosets, remain unclear. Anhedonic behavior describes the decreased ability to feel or experience pleasure^22^. The SPT is used to assess anhedonic behavior in models of depression, including those stress-induced, drug-induced, and genetically-induced^23–26^. Generally, the models display lower sucrose intake/preference, but this decline is recovered by chronic treatment with antidepressants. However, there are no SPT studies of marmosets. Other tests for motor activity (Activity) and food intake (FI) are already established, and there are standard methods routinely used for body weight (BW)^27,28^. In addition, the ES test is a test developed independently to examine food sorting behavior.

In the present study, a three-step work process was carried out to develop a new DSM-based research platform to determine depressive-like marmosets. First, several tests suitable for marmosets were designed and developed, and preliminary experiments were conducted to obtain basic data for each test. Next, the tests were combined to construct a test battery, and the criteria, order of implementation, and period were set. Finally, a usability confirmation experiment was conducted using drugs. Here, a drug that causes depression-like symptoms was administered to the marmosets, and changes were examined during the drug administration (ON) and non-administration (OFF) periods to confirm the drug’s efficacy. Depression-like marmosets were determined according to a simple and specific procedure (algorithm) based on patterns of change. The drug used was reserpine, which causes depression-like symptoms in laboratory animals and humans and has abundant human data ^29,30^. Using our new platform, we clarified how many of the nine drug-treated marmosets were depression-like marmosets and compared the incidence with previous human findings. This research platform, the “DSM Test Battery for Depressed Monkeys,” has the potential to establish new criteria for determining depression-like models and unify etiological testing and comparisons.

## Results

### Experiment-1: Duration and frequency of VEB by means of PhT

In experiment-1, a test was developed to measure the duration and frequency of peeping in response to curiosity while the test subject was in its home cage **(Fig. 1)**. We examined how the VEB of marmoset differed in response to different stimuli **(Fig. 2)**. The duration and frequency of VEB were investigated among five testing stimuli: a model insect, a model car, and three conspecifics (Conspecific 1, 2 and 3). A one-way repeated measures ANOVA showed that there were significant differences among the type of stimulus in both the duration (F _(4, 16)_ = 26.69, *p* < 0.001) and frequency (F _(4, 16)_ = 19.63, *p* < 0.001). By scrutinizing the data, the stimuli were divided into two main groups: object and conspecific. A multiple comparison Scheffe’s test showed significant differences between the object and conspecific groups (duration, *p* < 0.001; frequency, *p* < 0.001), indicating that the marmosets showed longer and more frequent VEB to the conspecifics than to the objects.

**Figure 1.**
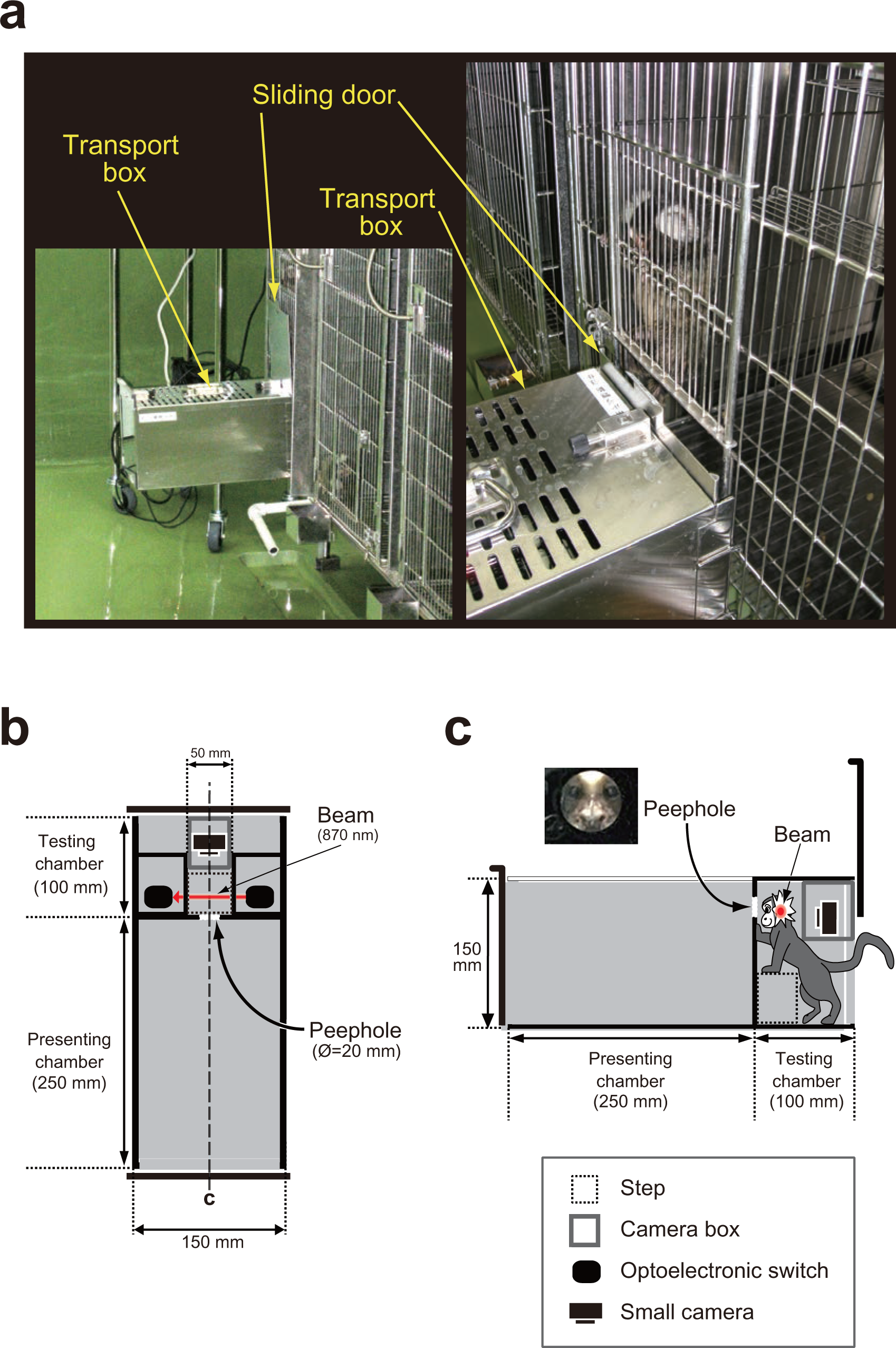
Experimental setup and apparatus of the peephole test (PhT) designed for common marmoset. (a) Left: the entire test apparatus during the experiment. Right: a subject marmoset and transport box attached to the subject’s home cage. (b) Ground plan of the apparatus. (c) Section plan on the center line drawn in (b). The inset photograph taken from the presenting chamber shows the peeping marmoset. The red arrow and spot indicate the infrared LED beam that detects the presence of the head.

**Figure 2.**
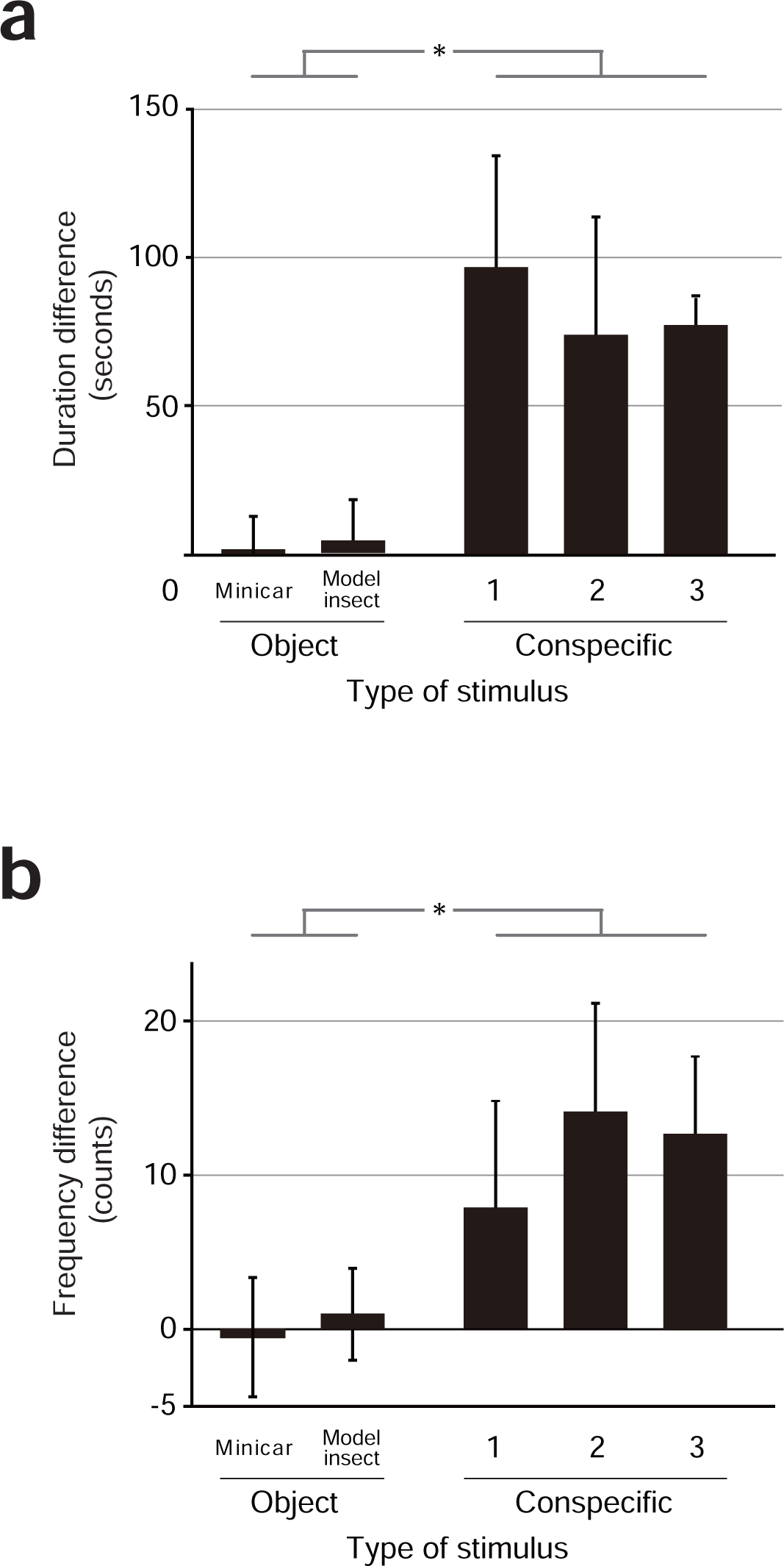
Characteristics of visual exploratory behavior (VEB) depending on the type of stimulus by means of the PhT. (a) Duration differences of VEB are shown for five different stimuli (two objects and three conspecifics). (b) Frequency differences of VEB are shown for five different stimuli. Data are expressed as the mean and SD. *p < 0.05, Scheffe’s contrast test (object vs. conspecific).

### Experiment-2: Sucrose concentration-response by means of SPT

The SPT is easily performed using a strong spring-loaded fixture, disposable bottle, and limiting the supply of water from other sources **(Fig. 3a)**. The animal intakes water from two bottles, one with sucrose solution and the other with tap water, over 24 days (4 days per concentration) **(Fig. 3b and c)**. A one-way repeated measures ANOVA showed significant differences in sucrose solution intake (F _(1.80, 10.79)_ = 49.79, *p* < 0.001), sucrose preference percentage (F _(1.56, 9.35)_ = 34.18, *p* < 0.001), and tap water intake (F _(5, 30)_ = 11.49, *p* < 0.001) depending on the sucrose concentration. Since a sucrose preference of 0% is assumed to be the anhedonic state, we compared sucrose solution intake and sucrose preference percentage with the 0% condition. A post hoc analysis revealed that sucrose solution intake was higher for sucrose concentrations of 0.375, 0.75, 1.5, and 3.0% compared with 0% (Shaffer method, *p* = 0.001-0.022), and the sucrose preference percentage was higher for 1.5 and 3.0% (Shaffer method; 1.5%, *p* = 0.017; 3.0%, *p* < 0.001). These results revealed that a sucrose concentration of more than 1.5% is needed to assess anhedonia in marmoset monkeys. Hence, we chose a sucrose concentration of 1.8% for the following experiments.

**Figure 3.**
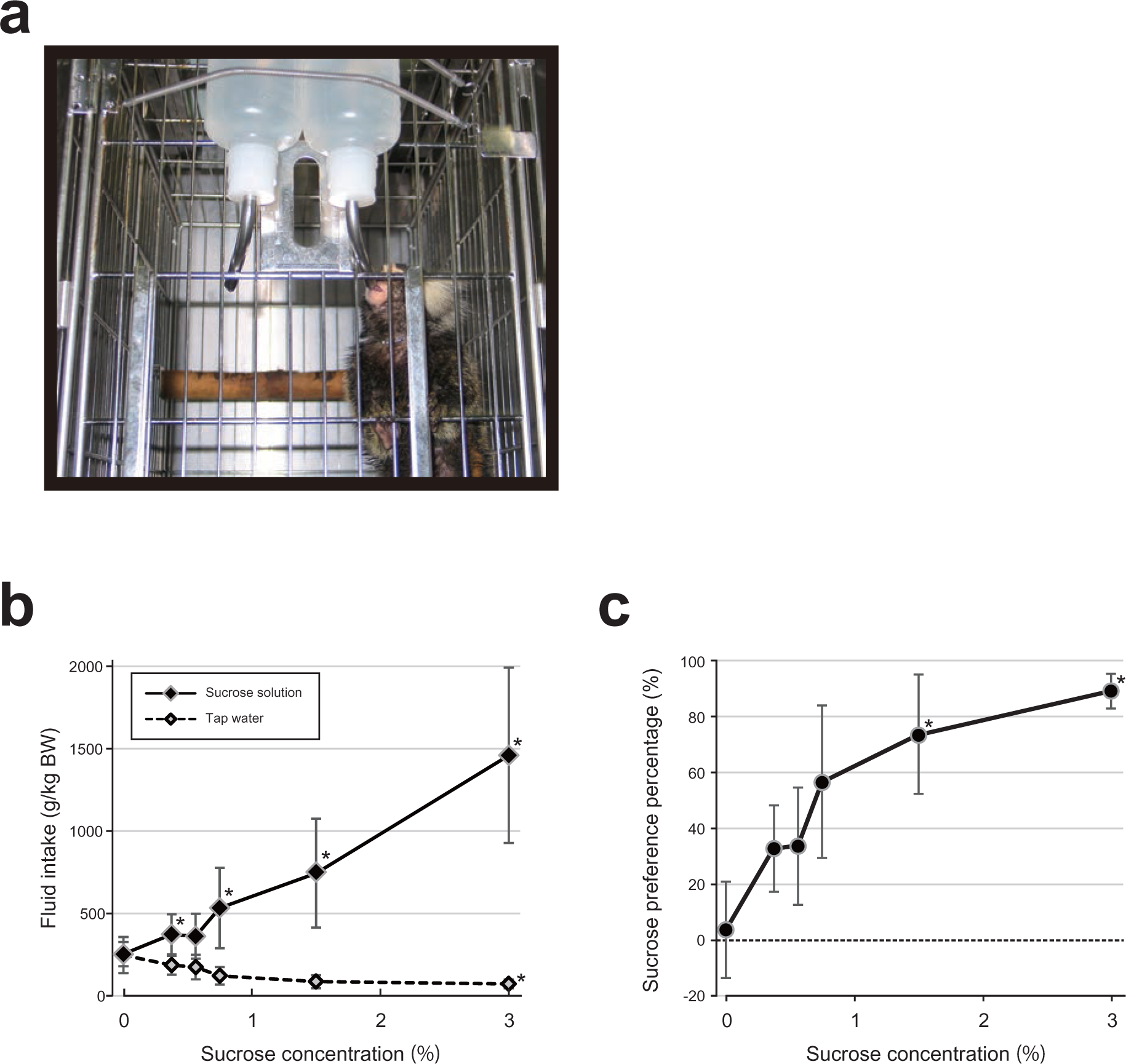
Setup of the sucrose preference test (SPT) and concentration-response curves. (a) A marmoset drinking from a bottle in its home cage. (b) Consumption of tap water and sucrose solutions against sucrose concentrations for four days. (c) Percentage of sucrose solution intake against concentration. Data are expressed as the mean and SD. *p < 0.05 compared to 0% sucrose (Shaffer’s test).

### Experiment-3: Preliminary experiment for ES test and FI

During the ES test, marmosets took pellets (food) from a feedbox and ate the food or dropped some into a mesh tray under their cages **(Fig. 4a and b)**. We successfully developed a measurement procedure consisting of five steps, which enabled us to simultaneously examine two indicators: the food drop index and intake **(Supplementary Fig. 1a)**. In this preliminary experiment of over 3 days, when 40.1 g of food was fed, the amount of FI was 12.5 g (SD 3.3 g), and the total amount of leftovers was 27.6 g (SD 3.5 g) **(Fig. 4c and Supplementary Fig. 1b)**. The amount of dropped leftover was 17.0 g (SD 7.5 g) resulting in a food drop index of 65% (SD 32.9%, range 19-100%) **(Fig. 4d and Supplementary Fig. 1b)**. Also, none of the 12 marmosets had a dropped leftover rate (food drop index) of 0%, indicating that they have the habit of dropping food if they are given enough food **(Fig. 4e**).

**Figure 4.**
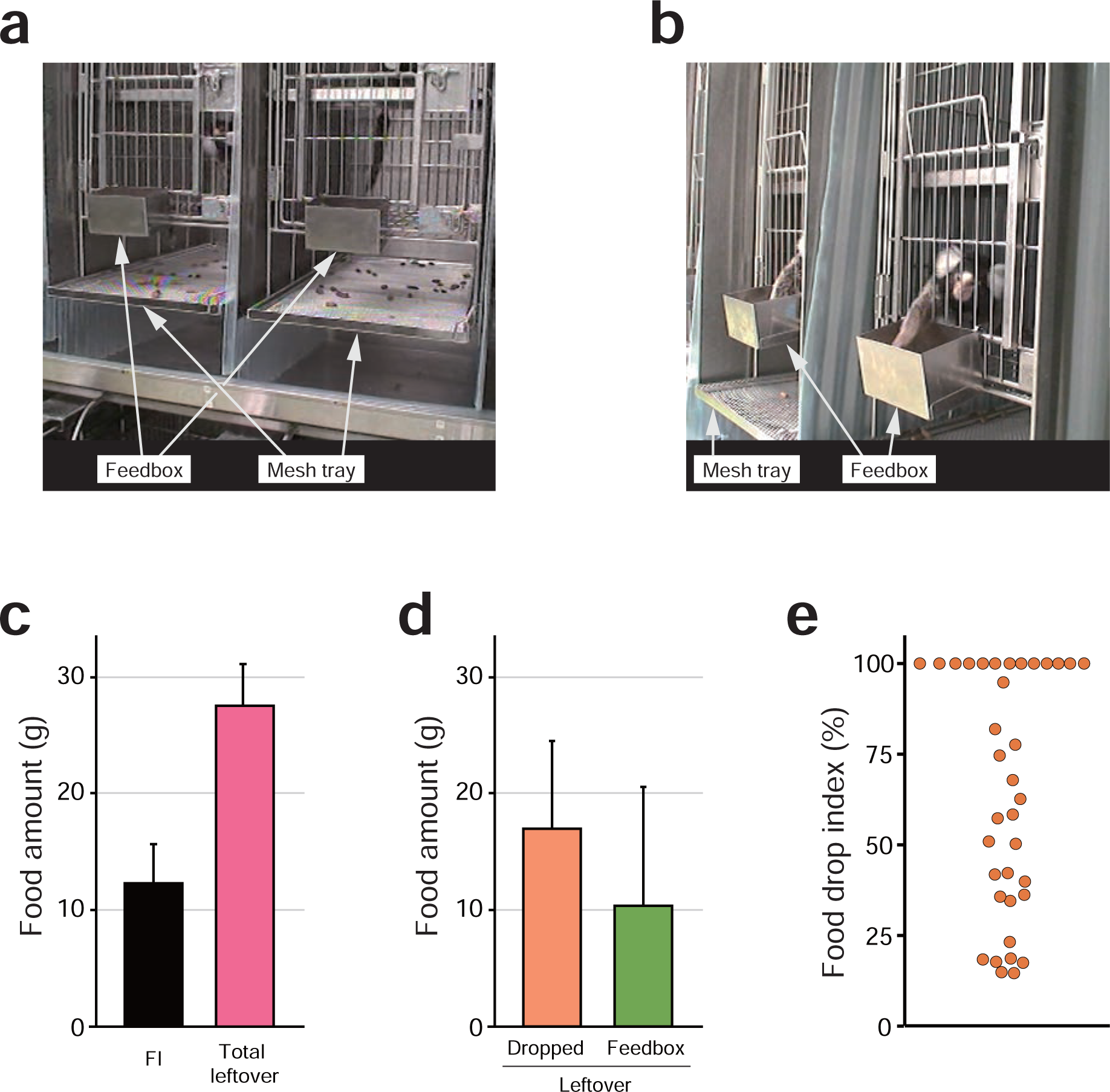
The eat style (ES) test and food intake (FI) in experiment-3. (a) A mesh tray placed under the floor of the home cage retrieves dropped food pellets. A feedbox is located on the front door of the cage, and marmosets have free access to the given food. (b) Two marmosets reaching out from the home cage and taking food from the feedbox. (c) The average FI and total leftover from a preliminary experiment. (d) The average dropped and feedbox leftovers from a preliminary experiment. Data are expressed as the mean and SD (n = 12). (e) The distribution of all 36 data points (12 animals, three-day trials each) for the food drop index.

### Experiment-4: Preliminary experiment for motor activity using an infrared sensor

Each marmoset’s hourly activity is shown in **Fig. 5**. Activity began in the morning, remained high during the day, and disappeared when the lights went out in the evening. The circadian rhythm of activity levels associated with changes in light and dark was consistent with previous reports about marmoset^27,31^.

**Figure 5.**
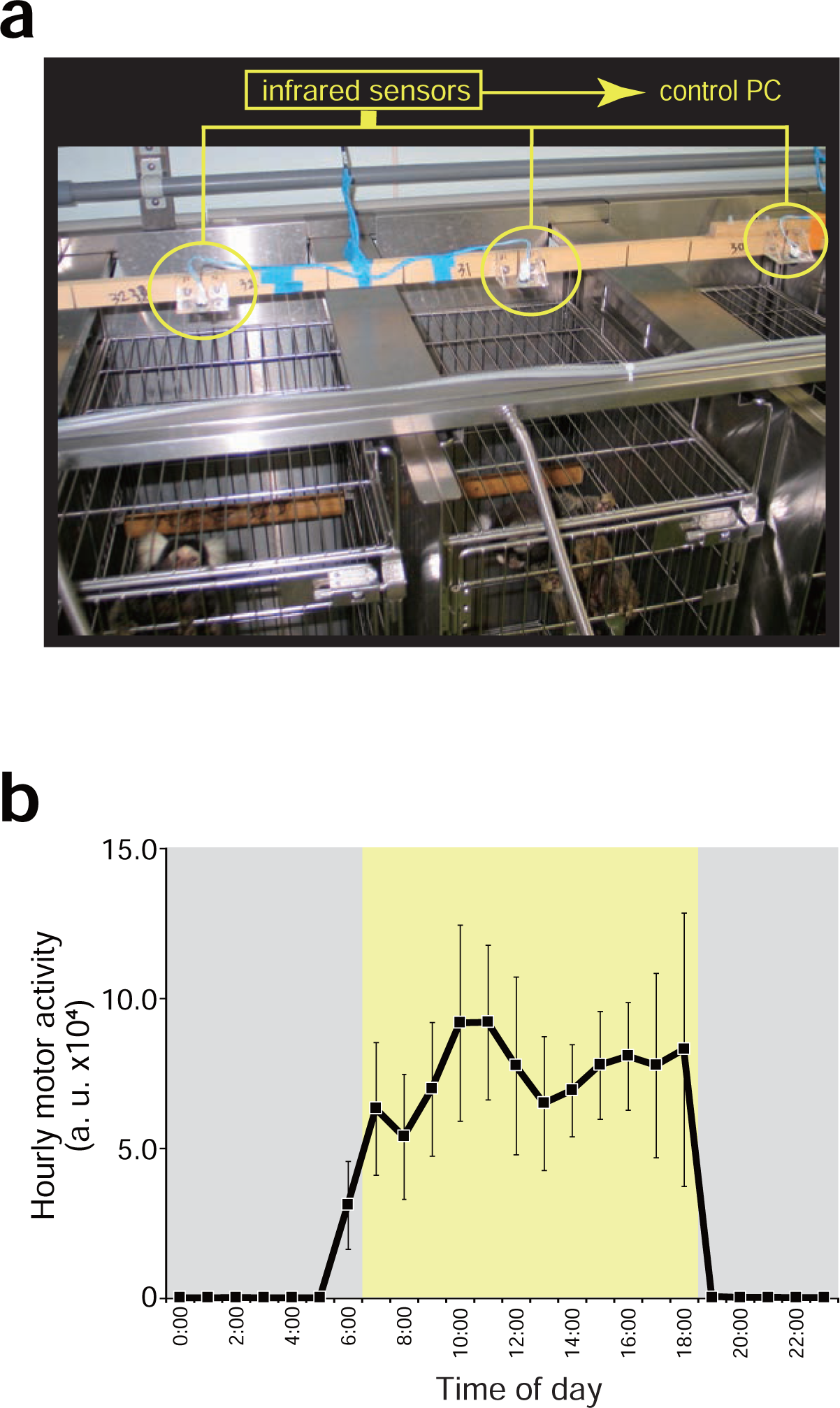
Measurement system and circadian rhythm of motor activity. (a) An infrared sensor is set above the ceiling of each home cage, and the acquired signal data is sent to a control computer in a separate room. (b) 24-hour observation of motor activity using the infrared sensor (n = 12). Black squares and bold lines indicate hourly activity acquired over the course of a week. The yellow area indicates the light period, and the grey areas indicate the dark period. Data are expressed as the mean and SD.

### Experiment-5: Usability confirmation experiment by a test battery consisting of six tests

To confirm how many marmosets were depression-like upon reserpine treatment, we identified individual marmosets showing depression-like symptoms using a test battery and specific judgment criteria. This battery consists of six tests: PhT, SPT, ES test, FI, BW, and Activity **(Fig. 6** and **Supplementary Table 1**). Three to six marmosets qualified as responders for each test (for details, see **Table 1** and **Supplementary Tables 2-8**). Three marmosets, Monkeys B, C and I, met the criteria that depression-like marmosets are responders in four or more tests, giving us an incidence of depression-like marmosets by reserpine treatment of 33%. This incidence rate resembles the reported 19% and 26% human clinical incidence^32,33^. Notably, Monkey C qualified as a responder for all six tests (**Fig. 7**), and Monkeys B and I qualified as responders for four tests (**Supplementary Fig. 2**). Plasma cortisol levels showed many marmosets qualified as responders, consistent with a previous rat study (**Supplementary Table 9**)^34^.

**Figure 6.**
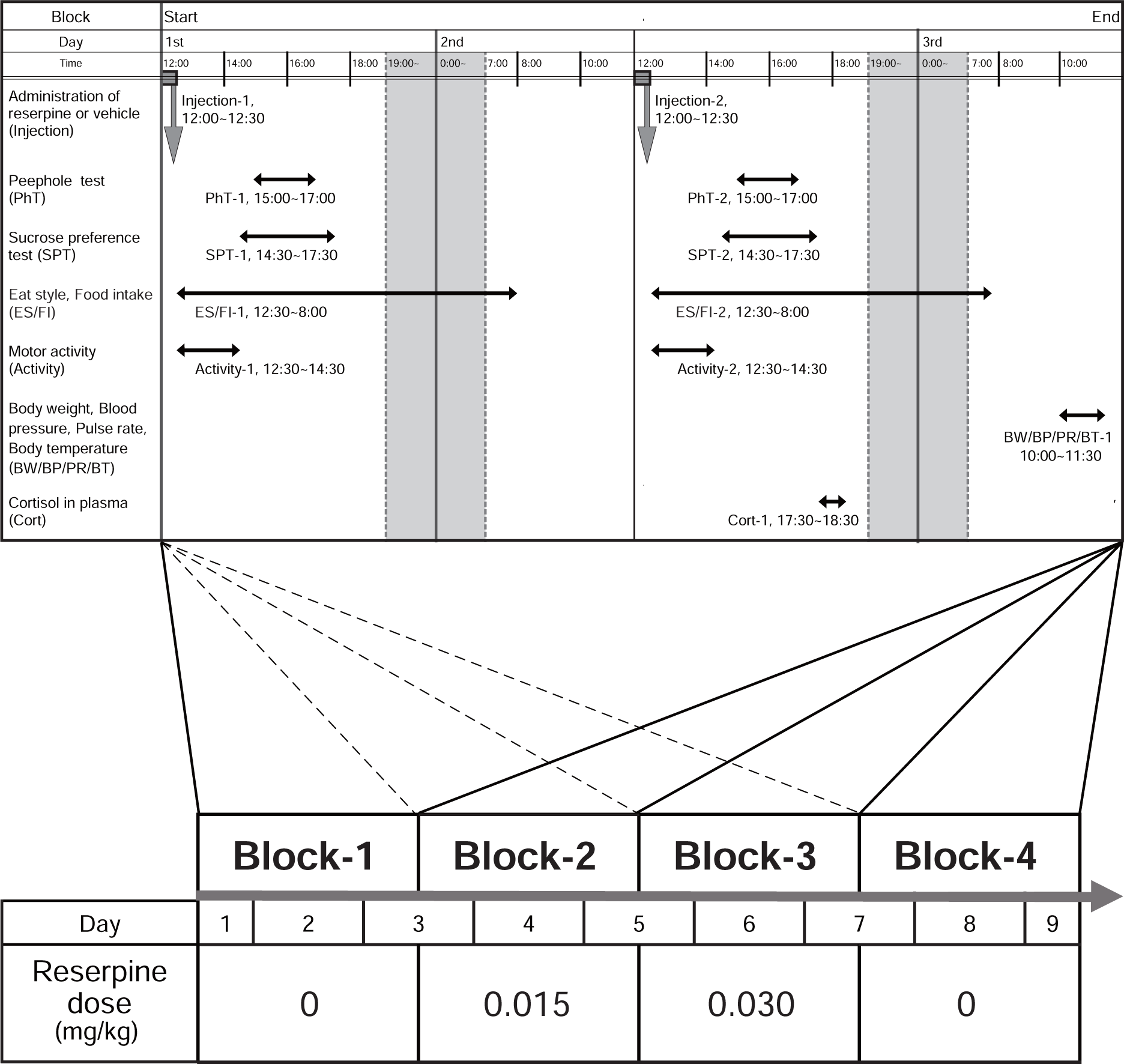
Experimental schedule for the depression-like marmoset in the test battery system in experiment-5. One block consists of 2 days, and 10 tests are conducted in a block. Continuous measurements for 8 days (four blocks; 2 days each) are performed, and reserpine-treated blocks (Block-2 and -3) are sandwiched between the two vehicle-treated blocks (Block-1 and -4). Most of the six tests are performed daily, and vital measurements and blood sampling for cortisol are performed once per block as additional reference data.

**Figure 7.**
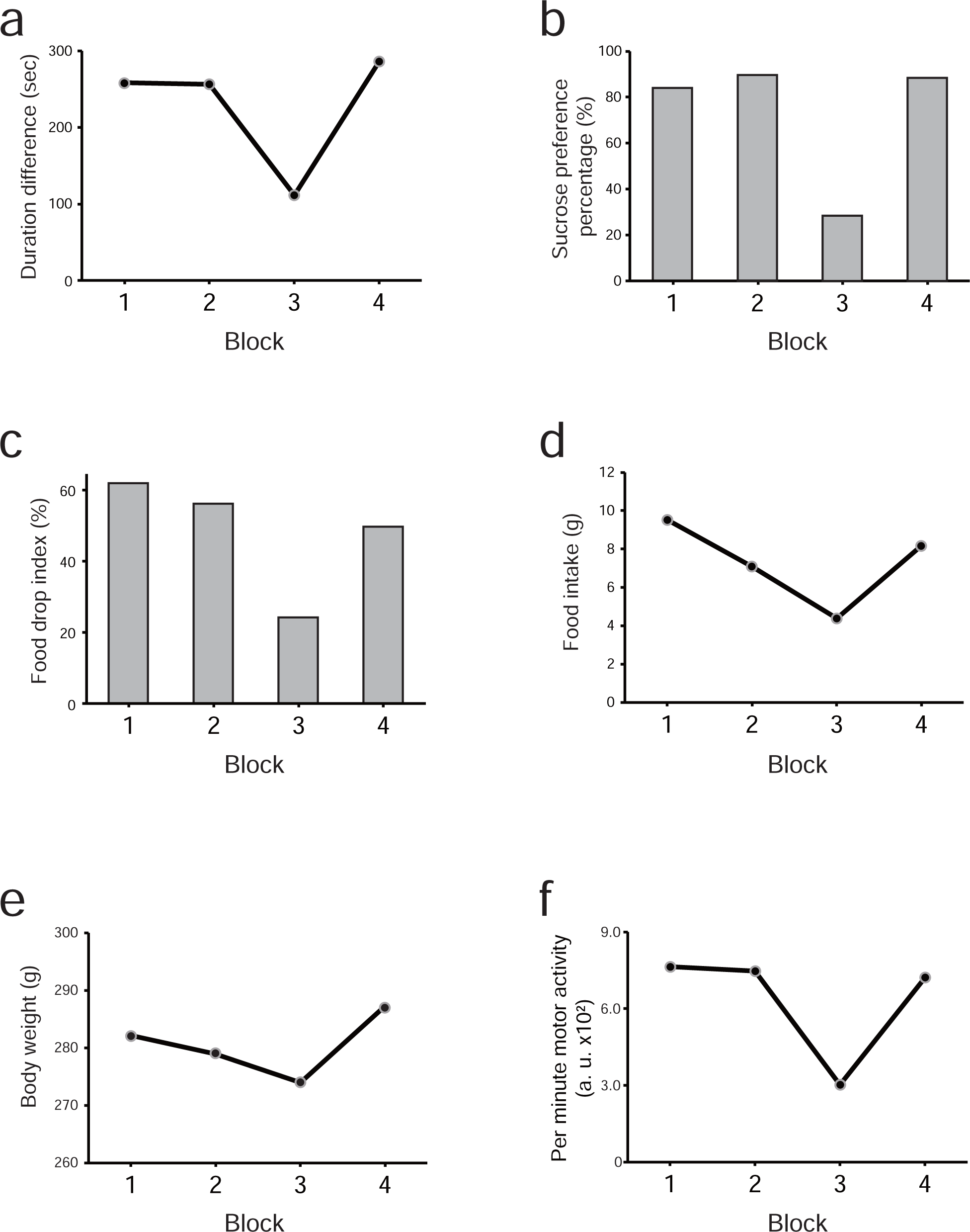
Depression-like symptoms of Monkey C in the six tests of the battery. Drug-induced V-shaped change patterns were observed in the (a) duration difference of VEB in the PhT, (b) sucrose preference percentage in SPT, (c) food drop index in the ES test, (d) food intake, (e) body weight, and (f) motor activity.

**Table 1.** Responder list for each test of the battery in experiment-5.

| Test name | Monkey name |  |  |  |  |  |  |  |  |
| --- | --- | --- | --- | --- | --- | --- | --- | --- | --- |
|  | A | B | C | D | E | F | G | H | I |
| Peephole test (PhT) | – | **R** | **R** | – | – | R | – | – | **R** |
| Sucrose preference test (SPT) | – | – | **R** | – | R | – | – | R | **R** |
| Eat style (ES) test | – | **R** | **R** | – | R | R | – | – | **R** |
| Food intake (FI) | R | **R** | **R** | – | R | – | – | – | **R** |
| Body weight (BW) | – | – | **R** | R | – | – | – | R | – |
| Activity | R | **R** | **R** | R | – | – | R | R | – |
| Number of tests qualified as responders | 2 | **4** | **6** | 2 | 3 | 2 | 1 | 3 | **4** |

“R” indicates responder marmosets who displayed a V-shaped pattern,
“–” indicates non-responders who did not display a V-shaped pattern.
Bold letters indicate depression-like marmosets. If in more than half of the tests (at least four tests) the subject qualified as a responder, then the individual was judged to be a depressed-like marmoset.

## Discussion

Here we report the “DSM Test Battery for Depressed Monkeys”, a new research platform for the study of depression using marmoset. We designed and developed several tests for marmosets and collected their basic data. We clarified in these preliminary experiments that VEB is efficiently induced when a conspecific is presented as the stimulus in the PhT and that sucrose water at a concentration of 1.5% or higher should be used to identify anhedonic behavior in the SPT. Next, we combined tests and set the criteria, order of implementation, and period for this platform. Finally, in a usability confirmation experiment using reserpine, the incidence rate of depression-like marmosets detected by this battery test was 33%, which is comparable with the human incidence^32,33^.

This battery covers four of the nine DSM diagnostic criteria (**Supplementary Table 1**). Among the remaining five, it is possible to add a new test for (4) abnormality of sleep and (8) diminished ability to the battery. The former can be measured using sleep polysomnography, and the latter can be examined using a cognitive function test^35,36^. Because the outstanding three criteria are based on the patient’s chief complaint and information from their families, they are difficult to introduce to the battery. Importantly, this battery is close to a clinical diagnosis in that the symptoms are evaluated for each and every individual, facilitating translational research.

Using the battery as a screening strategy for an exploratory study will enable the isolation and identification of individuals, etiology hypotheses, and the selection of antidepressants with certain characteristics. For example, since the depressive symptoms caused by reserpine do not appear in all individuals even at the same dose, it is generally assumed that there is a hidden regulator system determining the vulnerability or responsiveness in the background of the depression. In order to identify this system, it is necessary to efficiently obtain and use a depression model with high phenomenological similarity. Based on our finding, we therefore concluded that Monkey C should be bred over several generations. In this regard, the battery will play a critical role in selective breeding. Among primates closely related to humans, the marmoset is advantageous for selective breeding because of its earlier sexual maturation and higher fecundity. In addition, this strategy may selectively breed individuals who are not only highly responsive to reserpine but also vulnerable to depression.

As for the test battery system’s use to study etiology hypotheses, it has been reported that synaptic transmission and neuronal activity can be switched on and off by drug administration in DREADD (Designer Receptors Exclusively Activated by Designer Drug) and Tet-on systems using a virus vector. This capability allows for an analysis of V-shaped patterns using doxycycline or clozapine-N-oxide^37,38^.

Finally, regarding use of our battery in preclinical antidepressant screening, new antidepressant candidates have been screened mainly using rodent models. Moreover, the battery may be able to pick up other antidepressants that do not show positive effects in rodents. For example, the teratogenicity of thalidomide and the appearance of major signs of MPTP are observed in primates but not in rodents^39^. By setting a sufficiently long washout period between treatments, different drugs can be evaluated by the same assessment tool using an identical individual, resulting in more strict and effective comparisons of drugs.

The PhT revealed that VEB in marmosets has similar characteristics as rhesus monkeys. Thus, this test will be a useful tool not only for evaluating symptoms of depression but also for researching interpersonal relationships, fear, and the development of antidepressants and anxiolytics. The presentation of a conspecific individual has been reported to increase the frequency of VEB 2.7-fold compared with the absence of the individual^19^. In the present study, we found the mean frequency was 16 counts under the presence of a conspecific in the stimulus phase and 5 counts in the empty chamber condition in the baseline phase for a 3.2-fold change in experiment-1. The similarity of this fold change to that observed in rhesus monkeys supports the idea that the PhT is a useful tool for measuring VEB in marmosets. Also, the PhT is useful for objectively evaluating attachment, interest, and curiosity by focusing on VEB as well as research on social aversion and withdrawal, anxiety, and fear. For example, an ethologically based test using a taxidermized predator (e.g., cat, snake, or hawk) that is placed on a large arena was reported to study anxiolytic action in marmoset^40^. Similarly, it is possible to measure the duration and frequency of VEB only by replacing the stimulus in the chamber with a taxidermized predator in the PhT apparatus. Hence, in a smaller space, it is possible to examine the effects of psychotropic drugs such as anxiolytic drugs on behavior. On the other hand, we found no responder to the conspecific-induced frequency difference of VEB in experiment-5. Furthermore, the frequency in control block-1 (2.1 ± 5.0 counts, n = 9) was low compared with that in experiment-1 (11.5 ± 6.0 counts, n = 5), and a large number of marmosets showed values near zero or negative. In contrast, the duration difference had some responders, remained high and stable, and was significantly unaffected by the time course from experiment-1 to experiment-5. Thus, the frequency of VEB might be a labile indicator with an elusive property. In addition, the PhT in the next generation of the battery should use only conspecifics as the presenting stimulus.

We also reported, for the first time, the sucrose concentration-response curves of the SPT in marmoset, although the SPT has been reported in monkeys (rhesus and marmoset) and rodents. Although the response curve in the marmoset showed a concentration-dependent increase similar to that of the rhesus monkey, there was a difference in slope, with the marmoset showing a weaker relationship. Additionally, the curve has been studied in a maternal-deprivation depression model, with the consumption of sucrose solution reducing at 1.5% and 3.0% concentrations, consistent with our findings^41^. The sucrose concentration used in the present study was 1.8%, which is within these two concentrations. On the other hand, the sucrose concentration used in the SPT for rodents is typically 0.5–2%. If the concentration is too low, individuals will not show a preference for sweetness. Conversely, if it is too high, the sensitivity will be poor and it will be difficult to detect anhedonic behavior. Thus, 1–2% sucrose concentration might be optimal for investigating the anhedonic behavior of marmoset in the SPT.

The food drop index in the ES test is an essential test in depression research but is also underdeveloped. The food drop index is defined as the percentage of pellets thrown on the floor after being picked up once, nibbled, and checked for favorite flavor and softness and describes a series of actions referred to as food sorting behavior. In the present study, numerous pellets were observed under the animals’ home cages, and some of the pellets showed evidence of nibbling. This observation suggests that the marmosets engaged in sorting the pellets by grasping and evaluating their seasoning and texture. A decline in dropped foods is associated with symptoms such as diarrhea and disturbed fur. Therefore, monitoring the amount of dropped food serves as a crucial indicator in rearing practices for the early detection of poor health. In addition, since this behavior is a simple routine task and does not require a high level of concentration or thinking, it can be used to evaluate fatigue and loss of energy (symptom number “6”) in the DSM criteria **(Supplementary Table 1)**. The food drop index was high (65%), and experiment-3 revealed that healthy marmosets always dropped food selectively. Maintaining stable high levels in this food drop index is essential for disease modeling research that focuses on assessing reductions in the food drop index related to DSM symptom 6. Initially, it was hypothesized that the food drop index serves as the same test indicator as food intake. However, it was determined that the two indicators are different, as clear differences were observed between responders **(Table 1 and Supplementary Table 5-6)**. For example, Monkey F qualifies as a responder according to the food drop index, but not according to food intake. Additionally, Monkey F displays a clear V-shaped pattern in the food drop index, whereas its food intake continues to increase across blocks. By studying the food drop index with food intake, which can be measured simultaneously, we expect that the characteristics of the food index can be more clearly elucidated.

In conclusion, the present test battery is a valuable tool for assessing depression-like symptoms in common marmoset models, testing the etiology of depression and developing novel antidepressants. The test battery we propose has the potential to become a powerful translational tool to fill the gap between experimental animal models and clinical depression.

## Methods

### Animals

Experiments were performed on 14 marmoset monkeys (*C. jacchus*, aged 2.2 ± 0.5 years, 12 male and 2 female). Three additional male marmoset monkeys (conspecific, aged 2.4 ± 0.8 years) were used as stimuli in the PhT. All monkeys were housed indoors at room temperature (25–28 °C with 40–60% humidity). Light was provided from 07:00 to 19:00. Food pellets (CMS-1M, CLEA Japan Inc, Tokyo, Japan) softened with warm water and supplemented with honey, vitamins, and BIOFERMIN (intestinal regulators) were fed daily. Water was provided *ad libitum*, except in the SPT. The animals were maintained and handled in accordance with the “Guide for the Care and Use of Laboratory Animals” by the National Research Council in the United States and the guidelines of the National Institute of Neuroscience, National Centre of Neurology and Psychiatry in Japan. All experimental procedures were approved by the Animal Experimental Committee of the National Institute of Neuroscience.

### Peephole Test (PhT)

An acrylic plastic apparatus (150 x 350 x 150 mm) was placed into a stainless-steel transport cage (200 x 365 x 190 mm) that was attached to the subject’s home cage **(Fig. 1)**. The apparatus consisted of a testing chamber and a presenting chamber with a black wall in between. The wall had a 20 mm diameter circular window that serves as a peephole. The subject animal could enter the testing chamber and view a stimulus in the presenting chamber through the peephole. A transparent acrylic plate was set in the peephole to avoid direct physical contact between the subject individual and the stimulus individual. The subject, upon entering the test chamber, needed to duck under the camera box and step on the step box to peep inside the presenting chamber. The absence of extra space enabled detection of the subject’s head by an optoelectronic switch. The most important feature of the PhT device is its design, which requires the subject’s head to be positioned in a narrow space to peep through the hole. An optoelectronic switch (E3Z-T66, OMRON Co., Kyoto, Japan) and a small pinhole camera (KPC-S500P1, KT & C Co., Seoul, Korea) on the back of the hole detected the response of the subject. The optoelectronic switch consisted of an infrared emitter and receiver to detect if there was an obstruction in between. The camera monitored from behind the subject to simultaneously detect the subject’s presence. One test trial consisted of two distinct phases (stimulus and baseline phases), and only one stimulus a day was presented. The stimulus was present in the stimulus phase, and the presenting chamber was empty in the baseline phase. Two 300-s phases were performed in a trial; the baseline phase followed by a principal stimulus phase. The duration and frequency of VEB were recorded with an acquisition program written in LabView (National Instruments, Austin, TX) in another room. The duration and frequency of the baseline phase were subtracted from those of the stimulus phase throughout the analysis. Prior to the experiment, each marmoset was acclimated by exposure to the apparatus 13 times, with 20 min acclimation trials once per day. Five animals were subjected to experiment-1, and all measurements were made between 13:00 and 17:00. The PhT was conducted at intervals of less than or equal to 7 days, and two repeated measurements were performed for each of the five stimuli. Inanimate stimuli (object) consisted of a model car and a model insect, and three marmosets (Conspecific 1, 2 and 3) different from the subject marmosets were prepared for the animate stimuli **(Fig. 2)**. The data for two trials using each stimulus was averaged and used for further analysis.

In experiment-5, two PhT trials were performed for each block; a stimulus animal and an object were presented one by one as the stimulus in a block. A stimulus animal contained two marmosets (Conspecific 1 and 3), which were different from the subject marmosets, and only a model insect was used as the stimulus object. The presentation order was at random, and daily tests were performed between 15:00 and 17:00 **(Fig. 6)**.

### Sucrose Preference Test (SPT)

Two Falcon tubes with water nozzles each containing either tap water or sucrose solution were attached to the front side of the home cage **(Fig. 3)**. The automatic water supply was stopped during the test period. All fluid intake was normalized by body weight. The preference rate was calculated according to the following formula:

Sucrose preference percentage (%) = [(Sucrose solution intake - Tap water intake)/Total Intake) x 100].

In experiment-2, referring to a previous study with rhesus monkey, six concentrations (0, 0.375, 0.5625, 0.75, 1.5, and 3.0%) of sucrose solutions were set and presented at random^41^. Seven animals were used in this experiment. Each concentration was presented for 4 days, and the left and right presentation positions were counterbalanced and changed daily. Each daily test trial was performed between 14:30 on the first day and 14:00 on the second day (23.5 hours a day). Weighing, washing the bottle, and replacement of the fluids were performed between 14:00 and 14:30. The data over the 4 days for each concentration was summed and used for the analysis.

In experiment-5, two test trials were performed for each block, and the position of the 1.8% sucrose solution was changed daily. Daily tests were performed between 14:30 and 17:30 **(Fig. 6)**. Two test results per block were averaged and used for further analysis.

### Eat Style (ES) Test and Food Intake (FI)

Food was presented for 19.5 h from 12:30 to 8:00 the following day, and the leftover food was collected separately from two places, a feedbox and a mesh tray **(Fig. 4a and b)**. The feedbox leftovers were defined as leftovers collected from the feedbox, and the dropped leftovers were defined as leftovers dropped on the mesh tray suspended below the floor of the home cage. The sum of these two was referred to as total leftovers. Leftover food was collected and then dried at 65 °C for one day before being weighed **(Supplementary Fig. 1a)**. The FI was the value obtained by subtracting the total leftover food from the given 40 g of feed. The food drop index (%) was calculated according to the following formula:

Food drop index (%) = (Dropped leftovers (g)/Total leftovers (g)) x 100].

The food used was ordinary pellets given in daily life, and no other food or reward was given during the experiment. Basic daily data for three consecutive days was obtained in experiment-3, the preliminary experiment, in which 12 animals were subjected. The same procedure was used to obtain daily data in experiment-5, in which 9 animals were used.

### Motor Activity

We used home cages equipped with an infrared motion sensor (Panasonic Electric Works Co., Osaka, Japan) on the ceiling to continually record the motion of the animals in the animal rearing room **(Fig. 5)**. The system utilized a passive thermographic infrared sensor to monitor the heat emitted from the animals. The 3D localization of the heat source was monitored, and changes in this localization were recorded as movement. Animal data was monitored and recorded on a computer using a data acquisition program written in LabView.

Data for one week were obtained from 12 animals every 24 hours in preliminary experiment-4. In experiment-5, activity data were acquired between 12:30 and 14:30. The 60-min activity data between 40 and 100 minutes after the administration time of reserpine was utilized **(Fig. 6)**. The motor activity in this time zone was converted to a one-minute average to obtain average motor activity. Two trials for two days were averaged as block data of motor activity and used for the analysis.

### Drug Administration

Reserpine depletes monoamines in the brain by functionally inhibiting vesicular monoamine transporter 2^29^. Reserpine (Sigma, St. Louis, MO) was dissolved in glacial acetic acid and then diluted to 0.10 mg/mL with sterile water. Marmosets were injected intramuscularly with 0.015 and 0.030 mg/kg reserpine or vehicle once daily at 12:00-12:30 on the day of an experiment **(Fig. 6)**. All reserpine and vehicle solutions contained 3.75% glacial acetic acid.

### Responder and Depression-like Marmoset

Prior to experiment-5, a test battery of six principal tests was constructed by determining the experimental schedule (timing, sequence, frequency, etc.) and the criteria for responders and depressive-like marmosets **(Fig. 6)**. Individuals displaying depression-like symptoms were determined using the judgment criteria based on DSM in experiment-5 **(Supplementary Table 1)**. Ten tests were incorporated into the experimental schedule, ensuring that all were performed at least once in one block, and four blocks of continuous measurement for 8 days before and after administration of the drug were performed. Reserpine was administered in block-2 and block-3 with a dose-escalation method, and the dosing schedule was set to be sandwiched the reserpine blocks between block-1 and block-4, in which the vehicle was administered. Marmosets were evaluated individually between the four blocks. Responders to reserpine were qualified by whether the test data displayed a V-shaped pattern over the four blocks. Individuals whose measurement data satisfied the following conditions were qualified to be responders: Block-1 > Block-3, Block-4 > Block-3, and Block-2 > Block-3. Depression-like marmosets were defined as individuals who were responders in more than half (≥ four tests) of the six tests. This condition corresponds to the DSM criteria for depression, which requires more than half (≥ five) of the nine symptoms be present.

### Statistical Analysis

Experiment-1 and -2 data were analysed using a one-way repeated-measures analysis of variance (ANOVA) followed by Scheffe’s test^42,43^ (when comparing two groups, object vs. conspecific groups) or Shaffer’s^44^ (pairwise comparison) post hoc test. The data used were confirmed to be normally distributed by passing the Shapiro-Wilk normality test. Homoscedasticity assumptions were investigated before the use of the ANOVA tests. *p* < 0.05 was regarded as significant. All analyses were performed using SPSS software (Version 11.5, SPSS Inc., Chicago, IL).

## Data availability statement

The datasets generated during and/or analyzed during the current study are available from the corresponding author upon reasonable request.

## Supporting information

Supplemental Informations

## Acknowledgements

We greatly thank all members in the Animal Models for Human Disease, National Institute of Neuroscience, National Center for Neurology and Psychiatry, for their technical assistance. This study was funded by research grant 20B-10 for Nervous and Mental Disorders from the Ministry of Health, Labor and Welfare of Japan, “Development of nonhuman primate models for mental and nervous diseases,” and partly supported by a research grant from the Astellas Foundation for Research on Metabolic Disorders.

## Author contributions

All authors conceived the experiments. H.Y. and H.I. performed experiments. K.N. and H.I. acquired financial support. H.Y. analyzed the data. All authors wrote the manuscript, discussed the data and commented on the manuscript.

