## Supplemental Informations for "Development of a DSM test battery to determine depression-like marmoset"

**Supplementary Figure**

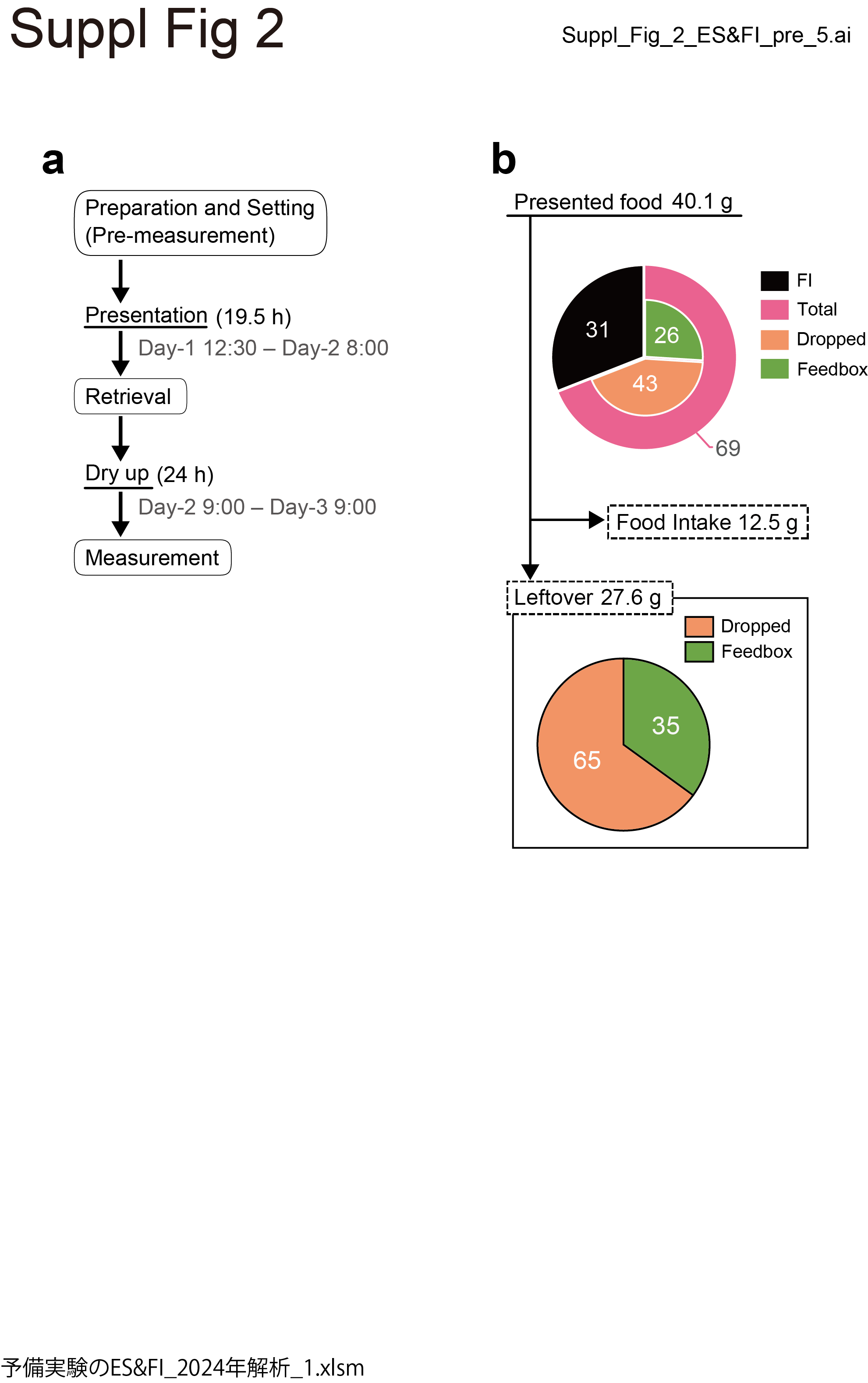

**Supplementary Figure 1**

Detailed experimental procedure and ratio of measured values for the ES test and FI. (a) The procedure consists of five steps and takes almost two days to complete the data acquisition for one day. The weight of the presented food, the feedbox leftovers, and the dropped leftovers are measured before and after the presentation. (b) 12 animals showed an average food drop index of 65% (SD, 33%) when enough food (40 g) was given.

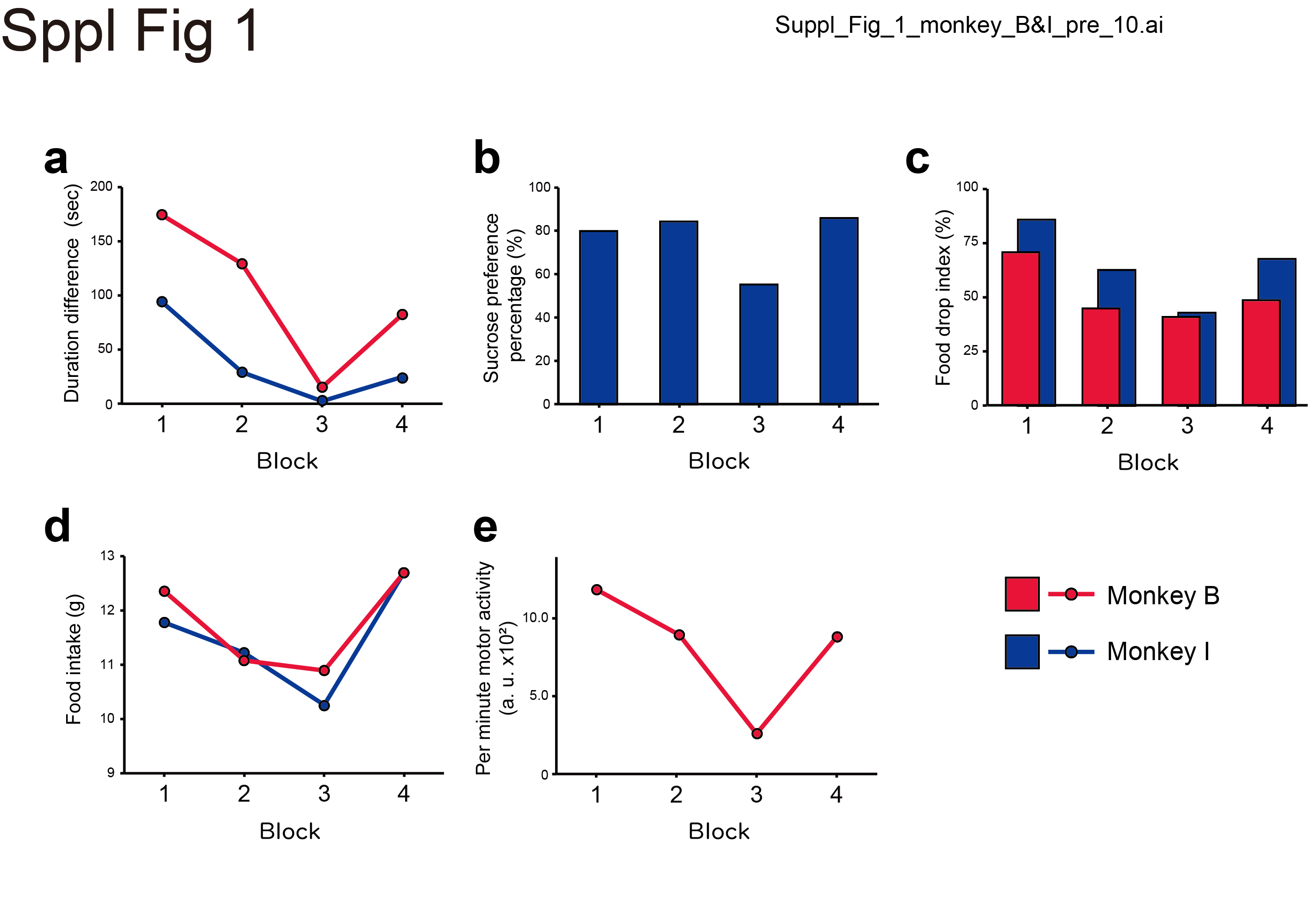

**Supplementary Figure 2**

Depression-like symptoms of Monkey B and I in five tests of the battery. Drug-induced V-shaped change patterns were observed in the (a) duration difference of VEB in the PhT, (b) sucrose preference percentage in SPT, (c) food drop index in the ES test, (d) food intake, and (e) motor activity.

**Supplementary Tables**

Supplementary Table 1. Six tests of the battery following DSM criteria for a major depressive episode.

|  | **Test name** | DSM criteria | Construct | Principal indicator |
| --- | --- | --- | --- | --- |
| 1 | **Peephole test (PhT)** | (2) Loss of interest or pleasure | Visual exploratory behavior (VEB) | Duration difference (seconds); Frequency difference (counts) |
| 2 | **Sucrose preference**  **test (SPT)** | (2) Loss of interest or pleasure | Anhedonic  behavior | Sucrose preference percentage (%) |
| 3 | **Eat style (ES) test** | (6) Fatigue or loss of energy | Food sorting behavior | Food drop index (%) |
| 4 | **Food intake (FI)** | (3) Abnormalities in body weight or appetite | Eating (appetitive) behavior | Food intake (g) |
| 5 | **Body weight (BW)** | (3) Abnormalities in body weight or appetite | Energy balance  (intake and consumption) | Body weight (g) |
| 6 | **Activity** | (5) Psychomotor agitation or retardation | Motor behavior | Motor activity (counts/min) |

The DSM diagnostic criteria symptom numbers are shown in parentheses, and the corresponding symptoms are underlined.

Supplementary Table 2. Conspecific-induced VEB in PhT

| Conspecific-induced VEB | | | | | | | | | | |
| --- | --- | --- | --- | --- | --- | --- | --- | --- | --- | --- |
| Duration difference | | | | | | | | | | |
|  | | Monkey name | | | | | | | | |
|  | | A | **B** | **C** | D | E | **F** | G | H | **I** |
|  | Block-1 | 19.9 | **176.5** | **258.5** | -34.4 | 159.7 | **32.4** | -23.2 | 196.7 | **94.7** |
|  | Block-2 | 46.3 | **130.7** | **256.5** | 53.4 | 247.4 | **15.1** | -2.7 | 160.3 | **29.5** |
|  | Block-3 | 56.0 | **15.7** | **111.2** | 46.9 | 238.5 | **0.0** | 0.0 | 0.0 | **2.5** |
|  | Block-4 | 19.6 | **83.6** | **286.7** | 23.2 | 202.0 | **25.5** | 10.2 | -17.9 | **25.4** |
| Frequency difference | | | | | | | | | | |
|  | | Monkey name | | | | | | | | |
|  | | A | B | C | D | E | F | G | H | I |
|  | Block-1 | 5 | 5 | 3 | -3 | 12 | 1 | -4 | 2 | -2 |
|  | Block-2 | 7 | 8 | -9 | 1 | 4 | 0 | -2 | -1 | -5 |
|  | Block-3 | 6 | 5 | 1 | -1 | 4 | 0 | 0 | 0 | 2 |
|  | Block-4 | 11 | 6 | 1 | 0 | 4 | 2 | -1 | -1 | 1 |

Bold letters indicate responders.

Supplementary Table 3. Object-induced VEB in PhT

| Object-induced VEB | | | | | | | | | | |
| --- | --- | --- | --- | --- | --- | --- | --- | --- | --- | --- |
| Duration difference | | | | | | | | | | |
|  | | Monkey name | | | | | | | | |
|  | | A | B | C | D | E | F | G | H | I |
|  | Block-1 | 1.1 | -4.6 | -18.4 | -7.2 | -1.1 | 0.2 | -1.4 | -3.4 | -39.2 |
|  | Block-2 | 0 | 0.7 | -20.3 | -4.6 | -25 | -4 | -3.2 | 14.7 | -7.6 |
|  | Block-3 | 0 | -0.7 | 0 | -5.3 | 39.6 | 0 | 0 | -21.8 | 0 |
|  | Block-4 | 0 | 0 | 3.5 | 7.5 | -28 | 0 | 2.6 | 14.3 | 13.8 |
| Frequency difference | | | | | | | | | | |
|  | | Monkey name | | | | | | | | |
|  | | A | B | C | D | E | F | G | H | I |
|  | Block-1 | 0 | -6 | -6 | -4 | 3 | -2 | 0 | -3 | -7 |
|  | Block-2 | 0 | 0 | -2 | -1 | -2 | -3 | -2 | -4 | -3 |
|  | Block-3 | 0 | -1 | 0 | -1 | 1 | 0 | 0 | -1 | 0 |
|  | Block-4 | 0 | 0 | -1 | 0 | 0 | 0 | 1 | 0 | 0 |

Supplementary Table 4. Sucrose preference percentage and sucrose solution intake in SPT

| Sucrose preference percentage (%) | | | | | | | | | | |
| --- | --- | --- | --- | --- | --- | --- | --- | --- | --- | --- |
|  | | Monkey name | | | | | | | | |
|  | | A | B | **C** | D | **E** | F | G | **H** | **I** |
|  | Block-1 | 88 | -20 | **84** | 2 | **97** | 67 | 84 | **96** | **80** |
|  | Block-2 | 92 | -60 | **90** | 87 | **94** | 93 | 77 | **93** | **85** |
|  | Block-3 | 90 | -44 | **28** | 95 | **87** | 69 | 83 | **90** | **55** |
|  | Block-4 | 95 | -80 | **88** | 95 | **96** | 76 | 77 | **93** | **86** |
| Sucrose solution intake (g/kg) | | | | | | | | | | |
|  | | Monkey name | | | | | | | | |
|  | | A | B | **C** | D | **E** | F | **G** | **H** | **I** |
|  | Block-1 | 57.8 | 4.3 | **42.4** | 49.3 | **91.2** | 38.5 | **45.0** | **110.7** | **44.8** |
|  | Block-2 | 60.3 | 3.3 | **39.5** | 77.3 | **76.3** | 68.9 | **43.6** | **102.7** | **42.0** |
|  | Block-3 | 54.5 | 5.3 | **14.0** | 87.5 | **68.2** | 44.7 | **38.9** | **85.6** | **28.7** |
|  | Block-4 | 48.8 | 1.9 | **46.3** | 99.6 | **85.1** | 48.6 | **41.2** | **89.8** | **49.8** |

Bold letters indicate responders.

Supplementary Table 5. Food drop index in ES test

| Food drop index (%) | | | | | | | | | | |
| --- | --- | --- | --- | --- | --- | --- | --- | --- | --- | --- |
|  | | Monkey name | | | | | | | | |
|  | | A | **B** | **C** | D | **E** | **F** | G | H | **I** |
|  | Block-1 | 100 | **71** | **62** | 83 | **75** | **56** | 22 | 31 | **86** |
|  | Block-2 | 100 | **45** | **56** | 77 | **100** | **53** | 32 | 12 | **63** |
|  | Block-3 | 100 | **40** | **24** | 79 | **72** | **28** | 31 | 36 | **44** |
|  | Block-4 | 100 | **49** | **50** | 77 | **100** | **49** | 19 | 9 | **68** |

Bold letters indicate responders.

Supplementary Table 6. Food intake

| Food intake (g) | | | | | | | | | | |
| --- | --- | --- | --- | --- | --- | --- | --- | --- | --- | --- |
|  | | Monkey name | | | | | | | | |
|  | | **A** | **B** | **C** | D | **E** | F | G | H | **I** |
|  | Block-1 | **19.4** | **12.3** | **9.5** | 17.3 | **13.6** | 12.0 | 11.7 | 9.7 | **11.8** |
|  | Block-2 | **16.6** | **11.1** | **7.1** | 13.0 | **12.9** | 12.1 | 9.3 | 9.9 | **11.2** |
|  | Block-3 | **16.2** | **10.9** | **4.4** | 14.2 | **12.2** | 15.3 | 11.7 | 9.7 | **10.2** |
|  | Block-4 | **16.4** | **12.7** | **8.2** | 14.6 | **12.9** | 18.3 | 14.4 | 11.4 | **12.7** |

Bold letters indicate responders.

Supplementary Table 7. Body weight in the test battery

| Body weight (g) | | | | | | | | | | |
| --- | --- | --- | --- | --- | --- | --- | --- | --- | --- | --- |
|  | | Monkey name | | | | | | | | |
|  | | A | B | **C** | **D*** | E | F | G | **H** | I |
|  | Block-1 | 352 | 340 | **282** | **338** | 309 | 448 | 493 | **305** | 368 |
|  | Block-2 | 347 | 338 | **279** | **338** | 305 | 446 | 480 | **303** | 369 |
|  | Block-3 | 352 | 338 | **274** | **345** | 309 | 452 | 481 | **301** | 368 |
|  | Block-4 | 356 | 338 | **287** | **342** | 306 | 455 | 490 | **304** | 370 |

Bold letters indicate responders.

*Monkey D, who displays an inverted V-shaped pattern, also qualified as a responder because DSM criteria include weight gain as well as weight loss.

Supplementary Table 8. Motor activity in the test battery

| Activity (counts/min) | | | | | | | | | | |
| --- | --- | --- | --- | --- | --- | --- | --- | --- | --- | --- |
|  | | Monkey name | | | | | | | | |
|  | | **A** | **B** | **C** | **D** | E | F | **G** | **H** | I |
|  | Block-1 | **886** | **1184** | **760** | **653** | 1533 | 1295 | **1011** | **1644** | 759 |
|  | Block-2 | **1045** | **893** | **744** | **840** | 548 | 478 | **378** | **590** | 124 |
|  | Block-3 | **534** | **260** | **304** | **452** | 1154 | 394 | **352** | **382** | 430 |
|  | Block-4 | **761** | **881** | **718** | **904** | 936 | 272 | **470** | **567** | 175 |

Bold letters indicate responders.

Supplementary Table 9. Plasma cortisol in the test battery

| Cortisol (μg/dL) | | | | | | | | | | |
| --- | --- | --- | --- | --- | --- | --- | --- | --- | --- | --- |
|  | | Monkey name | | | | | | | | |
|  | | **A** | **B** | **C** | **D** | **E** | F | **G** | **H** | **I** |
|  | Block-1 | **37** | **35** | **37** | **72** | **78** | 95 | **25** | **57** | **48** |
|  | Block-2 | **34** | **60** | **49** | **103** | **68** | 90 | **29** | **52** | **48** |
|  | Block-3 | **62** | **77** | **111** | **126** | **84** | 84 | **63** | **72** | **99** |
|  | Block-4 | **51** | **50** | **57** | **92** | **64** | 55 | **35** | **44** | **52** |

Bold letters indicate responders.

**Supplementary methods**

***Plasma cortisol***

Marmosets were set in a restraint device after capture rapidly, and blood samples were collected via the femoral vein with a syringe containing heparin. Blood was obtained under consciousness between 17:30-18:30. Samples were centrifuged for 10 min, and isolated plasma was immediately analysed. All samples were assayed in triplicate using AIA-360 (Tosoh Corporation, Tokyo, Japan), and the three measurements were averaged. Responders in cortisol were qualified according to an inverted V-shaped pattern.

***Body Weight***, ***Blood Pressure***, ***Pulse Rate, Body Temperature***

The physical condition of the monkey and serious side effects on the circulatory system were assessed using body weight and some vital signs. These data were obtained under consciousness between 10:00-11:30 at the end of each block. Blood pressure and pulse rate were recorded with a MK-2000 (Muromachi Kikai Co., Ltd., Tokyo, Japan) without heating and averaged from repeated measurements in triplicate. Body temperature was measured using an infrared non-contact thermometer (Thermofocus, Tecnimed Ltd., Varese, Italy). Then, the marmoset was moved to a transport cage to measure body weight before being returned to the home cage.
